# TRAIT2D: a Software for Quantitative Analysis of Single Particle Diffusion Data

**DOI:** 10.1101/2021.03.04.433888

**Authors:** Francesco Reina, John M. A. Wigg, Mariia Dmitrieva, Joёl Lefebvre, Jens Rittscher, Christian Eggeling

## Abstract

Single Particle Tracking (SPT) is one of the most widespread techniques to evaluate particle mobility in a variety of situations, such as in cellular and model membrane dynamics. The proposed TRAIT2D Python library is developed to provide object tracking, trajectory analysis and produce simulated datasets with graphical user interface. The tool allows advanced users to customise the analysis to their requirements.

Availability and implementation: the software has been coded in Python, and can be accessed from: https://github.com/Eggeling-Lab-Microscope-Software/TRAIT2D, or the pypi and condaforge repositories.

A comprehensive user guide is provided at https://eggeling-lab-microscope-software.github.io/TRAIT2D/.

## Introduction

Single Particle Tracking (SPT) is a method to quantify particle dynamics in a sample, such as lipids diffusing on biological membranes (1). This is indifferent to the microscopy technique adopted for detection, and in principle only requires the sample to be sparse enough so that the target particles be identifiable singularly.

Among the earliest example of SPT experiments, we can mention the testing of Einstein’s theory of Brownian motion (2). The introduction of electronic camera detection and innovative microscopy systems have made it possible to collect SPT data with framerates up to 50kHz (3–6).

While seemingly straightforward, the analysis of SPT data is not immune from potential pitfalls. Conventionally, trajectory analysis in diffusive systems is performed following Einstein’s description of Brownian motion (7) and similar models (8, 9). This theoretical derivation, however, does not take into account the localization error (10) and motion blurring (11). Overlooking these sources of error could lead to overestimate diffusion coefficients (12), or to erroneously detected subdiffusion (13).

Several toolboxes and algorithms have been produced to fulfill the need to identify and track single particles (14, 15, etc), and analyse their motion employing conventional Mean Squared Displacement analysis (16), Deep Learning (17, 18) and other methods (19, 20). However, while in many cases the source code is freely available, they are not platform-agnostic and fully open-source. On the other hand, tools available on free platforms, such as FIJI or ICY, are not easily customisable.

With the TRacking AnlysIs Toolkit, TRAIT2D, we aim to provide a fully customisable tool for particle tracking, simulation, and analysis. In TRAIT2D, intuitive graphical user interfaces facilitate users with little to no coding experience to develop their own analysis pipelines. A simple 2D tracking algorithm is provided to extract trajectories from SPT data. The source code allows more advanced users to customize data analysis. Furthermore, a track simulator is included as a validation tool of individual analysis pipelines. This allows the user to generate movies of diffusing particles at variable levels of signal-to-noise ratio, with a user-specified point spread function.

## Implementation

TRAIT2D is a cross-platform Python software package with graphical user interfaces (GUIs) to support SPT experiments. For simplicity, the software can be divided into three main sections: the tracker, the simulator and the data analyser, which can be used jointly (Figure 1). The details presented here are expanded in the Supplementary Notes.

**Fig. 1.**
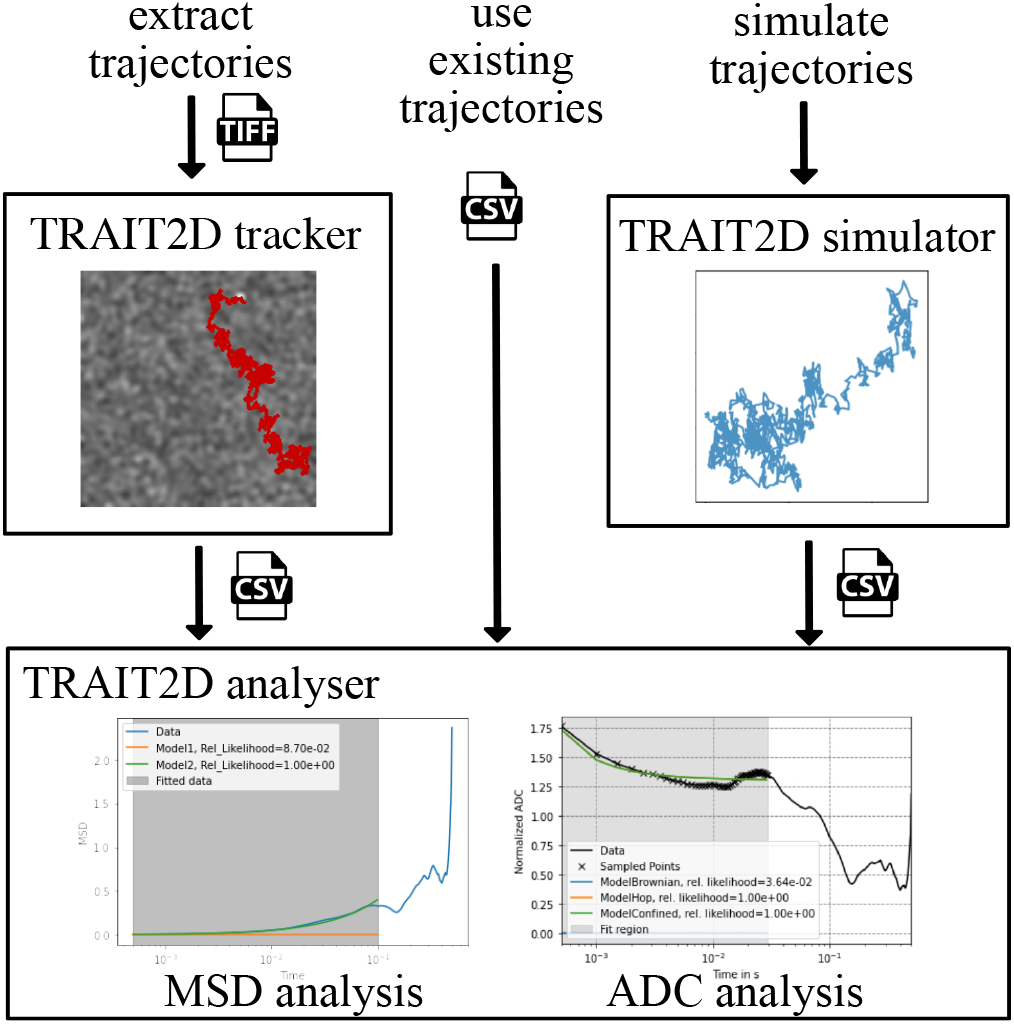
Workflow of the TRAIT2D software package. There are three possible avenues for the user to analyse their data: the user can import their timelapses, as tiff stacks, into the Tracker GUI and extract particle trajectories *(right)*, pre-determined tracks can be imported in the Analyser GUI from CSV files, with the importer function present in the Analyser GUI *center*, or control datasets can be simulated in the Simulator GUI, and analysed like experimental trajectories. Finally, the Analyser GUI will perform the analysis by following the Mean Squared Displacement (MSD) or the Apparent Diffusion Coefficient (ADC) pipelines (Supplementary Note 4).

The tracker extracts particle trajectories from a temporal image sequence. It allows a user to load and visualise TIFF timelapse stack.Once the image sequence is loaded, the parameters of the tracking can be set for detection, sub-pixel localisation and linking. The detection is implemented with Spot Enhancing Filter (SEF) (21), while sub-pixel localization is achieved using the radial symmetry center approach (22). Particle trajectory linking takes into account spatial and temporal distances between detections and exploits the Hungarian algorithm (23) for data association.

The simulator generates tracks in a virtual sandbox of arbitrary dimensions based on user-adjustable parameters, such as particle diffusion coefficient and time interval between localization. It is possible to simulate particles diffusing in simple Brownian diffusion and hop diffusion modes (24). Once the particle track has been generated, it is also possible to generate a complete movie by performing the convolution of the track with a point spread function, using varying levels of Signal-to-Noise ratio and pixelation. The last phenomenon has been implemented to model camera detectors usually employed for SPT. The purpose of this module is to provide computational controls and validation data sets for SPT detection and tracking algorithms.

Finally, the analyser allows the user to quantitatively evaluate tracks extracted from SPT movies following a Mean Squared Displacement (MSD) and Apparent Diffusion Coefficient (ADC) - based approach. Data analysis algorithms are accessible via the code or from a readily compilable GUI. Data can be obtained through the TRAIT2D-tracker, simulated by the simulator GUI or library, or imported from an externally generated comma-separated value (csv) file (Figure 1). The data analysis pipeline hereby employed is described in the Supplementary Materials (11, 12). The presence of a GUI for model fitting makes this procedure especially accessible and intuitive. The user can choose to fix or to put constraints on the fitting variables, as well as set the interval of data to be considered for the operation. It is also possible for the user to adopt alternative motion models which are not pre-compiled in the system (see documentation). The source code, together with exemplary Jupyter notebooks for easier understanding, can be retrieved at https://github.com/Eggeling-Lab-Microscope-Software/TRAIT2D. A comprehensive documentation can be found at https://eggeling-lab-microscope-software.github.io/TRAIT2D/.

## Conclusions

TRAIT2D is a Python-based, open source and easily deployable toolkit for the analysis and simulation of bidimensional SPT data. The combination of particle tracking, simulation, and trajectory analysis in the same package provides a number of benefits for the user. Another feature of this software are its user-friendliness for inexperienced users, owing to the presence of a clear graphical user interface, while retaining the potential for customisation for more advanced users.

## Acknowledgements

FR and CE would like to acknowledge Dr. B. Christoffer Lagerholm for his invaluable scientific advice. The authors would like to thank Dr. J. Keller-Findeisen for allowing the translation of the original algorithm for compartmentalised diffusion simulation.

