## Supplementary Notes for "TRAIT2D: a Software for Quantitative Analysis of Single Particle Diffusion Data"

### Supplementary information for the paper: TRAIT2D: a Software for Quantitative Analysis of Single Particle Diffusion Data

**Francesco Reina[1,2,✉], John M. A. Wigg[2], Mariia Dmitrieva[3], Joël Lefebvre[4], Jens Rittscher[3], and Christian Eggeling[1,2]**

[1]Leibniz-Institüt für Photonische Technologien e.V., Jena, Germany
[2]Institute of Applied Optics and Biophysics, Friedrich Schiller Universität, Jena, Germany
[3]Department of Engineering Science, University of Oxford, Oxford, UK
[4]Département d'informatique, Université du Québec à Montréal, Québec, Canada

#### Supplementary Note 1: Particle tracking

The particle tracking is developed to support a quantitative analysis module which requires extracted trajectories. We introduce a tool which allows setting tracking parameters, preview results and save final trajectories. The pre-processing of the image sequence is optional and can be applied prior to the tracking algorithm. The tracking is implemented as a two-step process. Firstly, the particles are detected using spot enhancing filter (SEF) (1) and sub-pixel localisation is estimated by the radial symmetry centre approach (2). Secondly, the detections are linked based on their spacial and temporal locations.
The current tool was developed for the Interferometric Scattering (iSCAT) imaging technique and the pre-processing step is built in accordance to the technique requirement. To distinguish the molecules of interest from the background, the image sequence is divided by a flat field. The flat field represents an undesired static background scattering. It is calculated by the pixel-wise temporal median filter over a set of frames (1000 frames in the current version). The background is calculated as a temporal average of the entire image sequence and subtracted from each frame. At last, the movie is normalized and brightness adjustment is applied to the image sequence (3). Although the pre-processing step is developed for the iSCAT it can also be employed for other techniques as a background subtraction tool.
The particle detection exploits Spot Enhancing Filter (SEF) to enhance the particles and reduce correlated noise in the image. SEF can be described by the convolution of the original image with a Laplacian-of-Gaussian (LoG) kernel and a thresholding to extract the spots. The threshold is defined by the average intensity of the image and its standard deviation, weighted by a constant $c$. This constant together with LoG parameter $\sigma$ can be defined by the user.
The local maximum of the image identifies a set of the detected particles in each frame. Further refinement of the particle coordinates (sub-pixel localisation) is implemented with the radial symmetry centre approach (2). It provides a faster execution time due to its non-iterative nature, while achieving high accuracy. The size of the region of interest for the sub-pixel localisation and the limitation for the detected peak size can be set manually.
The iSCAT imaging technique provides high temporal resolution. The small displacement of the particle between neighbouring frames allows reducing complexity of the linking algorithm and increase the computational speed. The linking approach focuses on strong spatial connection between frames. Hungarian combinatorial optimisation algorithm (4) is employed for the task of data association. The tolerance of the algorithm towards the special and temporal distance can be set by the parameters in the graphical-user interface. The assembled tracks can contain gaps in temporal domain due to the failed detection in one or few frames. These gaps are filled taking into account detections in the neighbouring frames and sub-pixel localisation. At the final stage, all the trajectories are filtered based on their length.

#### Supplementary Note 2: Track Simulation

The free (Brownian) diffusion was simulated by drawing a random walk on a two-dimensional virtual square space of side L, entered by the user. The particle is initialized in the centre of the square. The user also has the ability to select the value for the diffusion coefficient $D$ and the time step $dt$. The position of the particle is updated at each successive time step in the x and y direction separately:

$$x(t+dt) = x(t) + \sqrt{D*dt}*(\text{rand})$$
$$y(t+dt) = y(t) + \sqrt{D*dt}*(\text{rand})$$

This set of equations does not strictly respect the condition that $\overline{(x(t+dt)-x(t))^2 + (y(t+dt)-y(t))^2}$, since doing so would not account for the variability that happens in real simulations. Instead, the theoretical displacement $\sqrt{D*dt}$ is multiplied by a

random number ("rand") extracted from a normal distribution of mean$= 0$ and $\sigma = 1$.
The hop diffusion motion was simulated according to the code used for (5), kindly provided by Dr. J. Keller-Findeisen as a Matlab script, and transcribed into Python by the authors. First of all, a virtual surface of arbitrary dimension is split into an arbitrary number of compartments as a Voronoi diagram with random seed. To each of the compartments, a unique identifier is assigned. A particle is then generated in the centre of the plane, and its motion is simulated in the same way as the Brownian diffusion case. However, an additional condition is in place to simulate the hopping motion. A probability is assigned to the particle of jumping between one compartment and the other, called hopping probability ($HP$). Every time that the randomly generated displacement would move the particle to a different compartment, a random number between 0 and 1 is extracted. Should this number be less than the value of $HP$ initially fixed, the displacement calculation is repeated in order to keep the particle in the same original compartment.
Confined diffusion can easily be simulated by fixing $HP = 0$

#### Supplementary Note 3: Movie generation

We have created a simple movie simulator to obtain a synthetic movie from a list of detected or a simulated track. This simulator is inspired by a similar noisy holographic image simulator (6). Apart from a mandatory list of tracts to simulate, other parameters required for the simulation are the spatial ($r_{xy}$) and temporal ($r_t$) resolutions, the signal-to-noise ratio (SNR), the background signal level ($\mu_{bg}$), and the Gaussian noise level ($\sigma^2_{bg}$). Optional inputs are a point-spread function (PSF) stack, which can either be obtained experimentally or estimated with a model of the microscope, such as with DeconvolutionLab2 (7).
The simulator is initialised using the given tracks. The number of time frames, the minimum, and the maximum spot positions in $X$ and $Y$ are extracted from all tracks. These are used to initialise the simulation grid. The ($x_{min}$, $y_{min}$) and ($x_{max}$, $y_{max}$) positions define the simulation grid width and height. If the grid needs to be square, the min and max between ($x_{min}$,$y_{min}$) and ($x_{max}$, $y_{max}$) are computed respectively to define the grid shape. The simulation grid is discretised using the spatial resolution $r_{xy}$. For example, a track whose minimum position is (0,0), and maximum position is (10,10), and for a simulation resolution of $r_{xy} = 1$, will have a shape of $11 \times 11$, where each grid position represents a pixel of size $1 \times 1\,\mu m^2$. Furthermore, if the maximum time frame contained in a tract is 10, and if the temporal resolution is set to $r_t = 1$, the simulated movie will be of shape ($11 \times 11 \times 11$), where the first two dimensions represent the spatial dimension, and the last is the temporal dimension.
Once the simulation grid is initialised, we first add a background signal $B(x,y,t) \sim \mathcal{N}(\mu_{bg}, \sigma^2_{bg})$ everywhere. Each pixel background intensity follows a normal distribution of mean $\mu_b g$ and of variance $\sigma^2_{bg}$, and each simulated background pixels are independent and identically distributed random variables (i.i.d.). In other words, at this stage there is no spatial or temporal correlation between the pixels intensities. We are using the `util.random_noise` method from the `scikit-image` Python module (8).
Next, for each spot position in each track, we discretize the position in both the spatial $(x,y) \to (m_x, m_y)$ and temporal dimensions $(t) \to (m_t)$ using

$$m_i = \left[ \frac{x_i - x_{i,min}}{r_i} \right], \quad (1)$$

where $i \in \{x,y,t\}$ represents the $x$, $y$, and $t$ dimension of the spot positions respectively, $x_i$ is its position as recorded in the tract, $x_{i,min}$ is the minimum position within the tract, $r_i$ is the simulation resolution along the $i^{th}$ axis, and $[\cdot]$ represents the rounding operation to obtain discrete positions $m_i$ within the simulation grid coordinate framework. Each spot position is added iteratively to the simulated movie, and its intensity is given by the desired SNR for this simulation. At this stage, the simulated movie can be described by the equation

$$M(\mathbf{x}) = (1 + SNR) \sum_{i=1}^{N} \delta(\mathbf{x} - \mathbf{m}_i) + B(\mathbf{x}), \quad (2)$$

where $\delta(\mathbf{x} - \mathbf{m}_i)$ is a n-dimensional Dirac delta function, $\mathbf{x} = (x,y,t)$ is a $(3,1)$ vector representing the spatiotemporal position within the simulation grid, $\mathbf{m}_i = (m_{x,i}, m_{y,i}, m_{t,i})$ is the discretized vector position within the simulation grid of the $i^{th}$ spot, and $N$ is the total number of spots among all tracks to simulate. If a PSF is given as input, it will be applied at this stage with a convolution operator. If a 3D PSF stack is given, the central slice of the stack (corresponding to the focus position) will be used instead of the whole 3D stack for the simulation. The PSF can either be an experimentally acquired stack or a simulated volume, for example using the DeconvolutationLab2 with the microscope's configuration(7). To consider the boundary effects, we use zero padding. Thus, the simulated volume is

$$M(\mathbf{x}) = PSF(x,y) \otimes \left[ (1 + SNR) \sum_{i=1}^{N} \delta(\mathbf{x} - \mathbf{m}_i) + B(\mathbf{x}) \right], \quad (3)$$

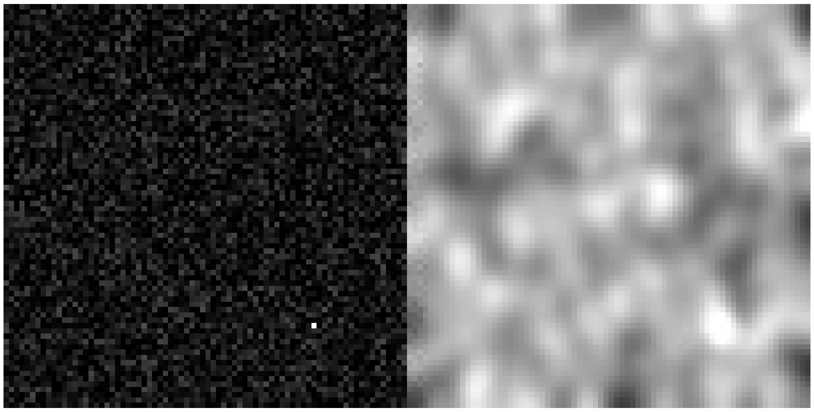

**Fig. 1.** Simulated iscat movie frame. (Left) Frame before and (Right) after the convolution by a PSF and the simulation of Poisson noise.

where $\otimes$ is the convolution product. Finally, i.i.d. Poisson noise is simulated for each pixel separately, using the value of $M(\mathbf{x})$ as the Poisson distribution parameter. The Poisson noise is simulated using the python module `scikit-image.util.random_noise`. Thus we obtain

$$M'(\mathbf{x}) \sim P(k;\lambda(\mathbf{x}) = M(\mathbf{x})), \tag{4}$$

where

$$P(k;\lambda) = \frac{\lambda e^{-\lambda}}{k!} \tag{5}$$

The simulated movie is exported as a tiff stack or an avi file. Figure 1 is a static example of a simulated movie frame using a single simulated track[1]. The image on the left is a movie frame before the PSF convolution and the Poisson noise simulation, and the image on the right is the final simulated movie frame after these operations.

#### Supplementary Note 4: Data Analysis Pipeline

In this section, we will briefly introduce the analysis methods implemented in the TRAIT-2D software package. The user-supplied SPT data must contain the coordinates of the particle at each moment in time, together with a track identifier in the case of multiple tracks. In this implementation, the analysis software only support analysis of uninterrupted tracks, that is, the localization happens at constant time intervals. There are two methods available to the user to analyse SPT trajectories that have been implemented in the software: the Mean Squared Displacement (MSD) analysis and the Apparent Diffusion Coefficient (ADC) analysis.
In the case of MSD analysis, the MSD is initially calculated according to the definition:

$$MSD(t_n) = \frac{1}{N-n-1} \sum_{i=1}^{N-n-1} |\underline{r}(t_{i+n}) - \underline{r}(t_n)|^2 \tag{6}$$

whereby $t_n = nt_0$ indicates the n-th time point from the initial time $t_0$, and $\underline{r}(t_n)$ indicates the two-dimensional position vector at time $t_n$. Notice that this notation, as well as the entire analysis pipeline hereby implemented, supposes that the time interval between localization is constant. According to Einstein's theory of Brownian Motion generalized for higher dimensionality (9, 10), this quantity is linked to the Diffusion Coefficient $D$ and time by the formula:

$$MSD(t_n) = 2dDt_n \tag{7}$$

in which n indicates the dimensionality of the system. In the rest of the section, we will assume $d = 2$ to simplify the formalism. In order to account for experimental finite localization precision, it is necessary to introduce an additive term to 7 (11):

$$MSD(t_n) = 4D(t_n)t_n + 2\delta^2_{x^2+y^2}. \tag{8}$$

The above formulation assumes that the localization error in the x and y directions is identical. This formula has been generalized to include the possibility of anomalous subdiffusion (12):

[1] A video version of this an animated version of this movie is available in this project Github repository

$$MSD(t_n) = 4D(t_n)t_n^{\alpha} + 2\delta^2_{x^2+y^2}. \quad \textbf{(9)}$$

While the user can, of course, fix the localization error value according to their chosen metric, it is also possible to leave it as a floating parameter. This is theoretically preferable from the point of view that the detection of moving particles is by necessity less precise than that of immobile particles, which are often used as a reference to estimate the localization precision of a microscopy system. The effect of motion blurring, to which this loss in precision is ascribable, is represented by a negative term by the form (13):

$$-8DRt_{lag} \quad \textbf{(10)}$$

where $R$ is a term which depends on the illumination mode of the chosen microscopy technique, and obeys $0 \leq R \leq 1/4$. It is especially interesting to notice how this term is dependent not on the time variable, but rather to the time resolution of the measurement. As a consequence of this, non-blur-corrected MSD data may suffer from considerable distortion at short time intervals, which coincidentally are the ones that are considered most important in determining the kind of motion the particle undergoes (14, 15). When this term is included in 8 and 8, it gives rise to the formulas:

$$MSD(t_n) = 4D(t_n)t_n + 2\delta^2_{x^2+y^2} - 8DRt_0 \quad \textbf{(11)}$$

and

$$MSD(t_n) = 4D(t_n)t_n^{\alpha} + 2\delta^2_{x^2+y^2} - 8DRt_0 \quad \textbf{(12)}$$

which are ultimately used to perform curve fitting.

The ADC analysis differs from the MSD analysis mainly for the central role given to the diffusion coefficient compared to Mean Squared Displacement. By adopting the convention

$$D_{app}(t_n) = \frac{MSD(t_n)}{4t_n\left(1 - \frac{2R}{n}\right)} \quad \textbf{(13)}$$

it is possible to switch to a diffusion coefficient-based representation. The nomenclature "apparent" is adopted to stress how this quantity is still affected by the localization error and not the absolute value of the diffusion coefficient.

Dividing both terms of 11 by $4t_n\left(1 - \frac{2R}{n}\right)$, and with some elementary operations, it is possible to derive:

$$D_{app}(t_n) = D(t_n) + \frac{2\delta^2_{x^2+y^2}}{4t_n\left(1 - \frac{2R}{n}\right)}. \quad \textbf{(14)}$$

By substituting in place of $D(t_n)$ the appropriate expression for the diffusion coefficient for a specific model, it is therefore possible to obtain a complete model to fit to the apparent diffusion coefficient data, obtained according to 13. In principle, it is possible to leave the localization error $\delta^2_{x^2+y^2}$ as a free-floating parameter in the fitting operation, to better estimate the localization error for moving particles.

The software package hereby presented comes with three built-in diffusion models (15, 16), corresponding to free (Brownian) diffusion

$$D(t_n) = D_M, \quad \textbf{(15)}$$

confined diffusion

$$D(t_n) = D_{\mu}\left(\frac{\tau}{t_n}\right)\left(1 - e^{-\frac{\tau}{t_n}}\right), \quad \textbf{(16)}$$

and compartmentalized diffusion, which is the combination of the above two models

$$D(t_n) = D_M + D_{\mu}\left(\frac{\tau}{t_n}\right)\left(1 - e^{-\frac{\tau}{t_n}}\right). \quad \textbf{(17)}$$

The nomenclature for these models follows the convention that $D_M$ is a constant diffusion coefficient, corresponding to theoretical Brownian diffusion, and that $\tau$ is the characteristic residence time in the confinement zones described by the confined diffusion model. $D_{\mu}$ is the microscopic diffusion coefficient observed in the aforementioned confinement zones, which can also be expressed as

$$D_{\mu} = \frac{L^2}{12\tau} \quad \textbf{(18)}$$

where $L$ is the average confinement zone size (17). We add that although the code we hereby describe has three built-in models, chosen according to the authors specific research interests, a documented procedure is in place to define a new model, according to the end user-specific wishes.
The selection of the best model to describe any given track is done by statistical means. Each of the models is fitted to the $D_{app}$ data, and the Bayesian Information Criterion (BIC) for each model is calculated according to the formula (18):c

$$\mathrm{BIC} = k\ \ln(n) + n\ \ln(RSS/n) \quad \textbf{(19)}$$

in which n is the number of data points the model is fitted to, the RSS is the residual sum of squares, and k is the number of degrees of freedom, i.e. the free parameters, of the fit. The most adequate model to describe the diffusion motion is then the one with the lowest value of BIC.
